# EEG-Beats: Automated analysis of heart rate variability (HVR) from EEG-EKG

**DOI:** 10.1101/2020.07.21.211862

**Authors:** Supakjeera Thanapaisal, Sabrina Mosher, Brenda Trejo, Kay Robbins

**Affiliations:** Department of Computer Science, University of Texas at San Antonio, San Antonio, TX, USA

**Keywords:** heart rate variability, human behavior, physiological indicator, big data, EEG, EKG, artifact

## Abstract

Heart rate variability (HRV), the variation of the period between consecutive heartbeats, is an established tool for assessing physiological indicators such as stress and fatigue. In non-clinical settings, HRV is often computed from signals acquired using wearable devices that are susceptible to strong artifacts. In EEG (electroencephalography) experiments, these devices must be synchronized with the EEG and typically provide intermittent interbeat interval information based on proprietary artifact-removal algorithms. This paper describes an automated algorithm that uses the output of an EEG sensor mounted on a subject’s chest to accurately detect interbeat intervals and to calculate time-varying metrics. The algorithm is designed for raw signals and is robust to artifacts, resulting in fine-grained capture of HRV that is synchronized with the EEG. An open-source MATLAB toolbox (EEG-Beats) is available to calculate interbeat intervals and many standard HRV time and frequency indicators. EEG-Beats is designed to run in a completely automated fashion on an entire study without manual intervention. The paper applies EEG-Beats to EKG signals measured with an EEG sensor in a large longitudinal study (17 subjects, 6 tasks, 854 datasets). The toolbox is available at https://github.com/VisLab/EEG-Beats.

## 1 Introduction

Heart rate variability is an indicator of many physiological and behavioral factors (Acharya et al., 2006) such as stress (Kim et al., 2018), fatigue (Vicente et al., 2016), and performance (Spangler et al., 2020) in normal subjects (Nunan et al., 2010) (Shaffer & Ginsberg, 2017). HRV measures can be extracted from signals generated from a variety of sensor types including electrocardiogram (EKG), electroencephalography (EEG), photoplethysmogram (PPG), and blood pressure monitors (ABP). HRV measures have been standardized (Task Force of the European Society of Cardiology and the North American Society of Pacing and Electrophysiology, 1996). Many open-source and proprietary tools have been developed for detecting peaks in cardiac signals and computing these measures.

The benchmark study by Vest et al. (Vest et al., 2018) compares five HRV toolboxes: PhysioNet HRV Toolkit (Goldberger et al., 2000), Kubios (Tarvainen et al., 2014), the Kaplan toolbox (Kaplan & Staffin, 1998), HRVTool (Vollmer, 2019), and the PhysioNet Cardiovascular Signal Toolbox (Vest et al., 2017). The benchmarks established that small differences in implementation can cause large variations in results. Vest et al. emphasize the importance of open algorithms with well-documented parameter settings and caution against using default parameters when analyzing raw EKG. Areas of particular concern include preprocessing, signal quality assessment for noisy segment removal, and detection of arrhythmias.

Traditionally, the focus of EKG in EEG experiments has been the removal of cardiac interference (Tamburro et al., 2019). Our interest in HRV arose from the possibility of using HRV indicators secondary measures of subject physiological state during EEG experiments. By placing a single EEG sensor on the chest, it is possible to extract HRV as a matter of routine during EEG data analysis. However, our initial efforts to analyze a large EEG/EKG study using several available toolboxes including Kubios and the PhysioNet Cardiovascular Signal Toolbox (PNC) was unsuccessful because peak detection often failed without extensive manual intervention.

In addition to large variations in both baseline signal levels and peak amplitudes, EEG signals also tend to have large signal bursts due to muscle activity and loose detectors. To address the difficulty of consistent large-scale extraction of HRV measures from EEG, the approach proposed in this paper uses a top-down, divide-and-conquer strategy to peak-finding in contrast to typical sliding window approaches such as Pan-Tompkins (Pan & Tompkins, 1985). While straightforward and robust to typical EEG artifacts, but the extraction method is applicable only to normal cardiac signals. The method detects positions of the peaks corresponding to heart beats and but not QRS complexes or other feature information such as arterial fibrillation or widespread arrhythmias.

This paper describes the HRV algorithm and an open-source implementation in a MATLAB toolbox. The toolbox, EEG-Beats, extracts inter-beat intervals with associated HRV measures and includes utilities for statistical analysis and visualization. EEG-Beats can be run in automated fashion on an entire EEG study. The results are validated by comparison with PNC, and examples of how the toolbox might be used are provided.

## 2 Methods and Materials

An *RR interval* is the distance between successive peaks without regard to the shape of the associated QRS complexes. *Normal RR intervals*, referred to in the literature as *NN intervals*, are not distinguished from other RR intervals in this paper. This section describes the automated EEG-Beats processing algorithms for extracting interbeat (RR) intervals, calculating HRV indicators from RR intervals, and analyzing the relationships of the extracted indicators. All explanations, computations, and examples in this paper use the EEG-Beats default parameter values. When values are given, the names of the corresponding settable parameters in parentheses appear in parentheses. EEG-Beats uses the statistics of an entire signal to determine peaks and should not be applied in an online fashion. Signals are recommended to be at least 5 minutes in duration.

In this paper, the central tendency is generally indicated by the median and distribution characteristics by the interquartile range (IQR). Thresholds are based on the *robust standard deviation*, defined as 1.4826 times the median absolute deviation from the median. The *robust outlier criteria* specifies that points that are 1.5 (*maxWhisker*) × IQR outside the first and third quartiles (mid quartiles) of a distribution are considered to be outliers.

### 2.1 Signal preparation

EEG-Beats downsamples the signal to 128 Hz (*srate*) and applies a [3, 20] Hz (*filterHz*) finite impulse response (FIR) filter to the EKG signal. The 3 Hz high-pass filter effectively removes trend, while the 20 Hz low-pass filter removes much of the high-frequency noise. The algorithm then subtracts the median signal and truncates the amplitude so that the signal is within 15 (*truncateThreshold*) robust standard deviations of the median signal. Truncation eliminates extreme signal spikes, which in the case of a loose detector can be thousands of times larger than the normal signal for EEG sensors.

### 2.2 Heartbeat detection

EKG acquired using EEG sensors has two distinctive characteristics that make setting of thresholds problematic for traditional heartbeat algorithms. First, the signal can undergo spikes in amplitude thousands of times larger than the base signal when a sensor makes poor contact. Second, the amplitude and signal directions can vary over time, as illustrated by the signal clip shown in Fig. 1.

**Fig. 1.**
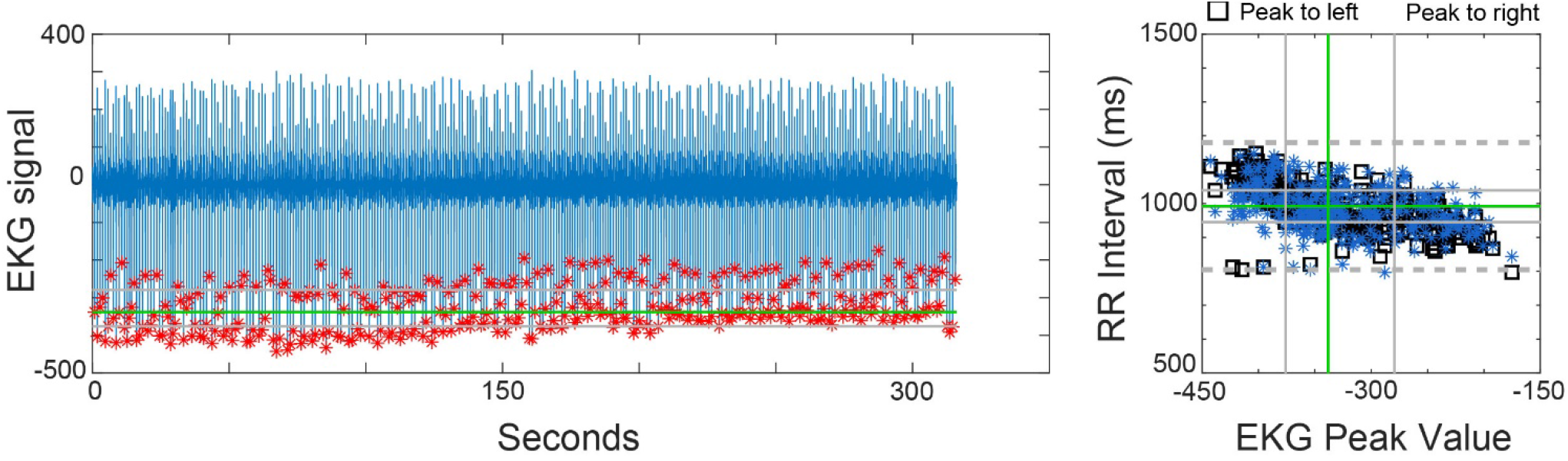
Graphical output produced by EEG-Beats. Left: EKG signal acquired using an EEG sensor (blue) for Dataset 1 (Subject S01) of the NCTU RWN_VDE study (Lin et al., 2018) as described below. Red asterisks show peaks identified by EEG-Beats. Right: EKG peak amplitude versus RR interval size for the peaks on the left (black squares) and right (blue asterisks) ends of each RR interval, respectively. Green lines mark medians and gray lines mark the interquartile ranges in each graph. Dashed gray lines on the right graph indicate the *robust outlier limits* as defined above. This dataset had no outlier peak amplitudes, so no gray vertical dashed lines appear on the right graph. The vertical axes for the right graph are always fixed to be the range [*rrMinMs, rrMaxMs*].

EEG-Beats determines the peak locations of the heart beats by first orienting the signal so that the peaks are always in the positive direction and troughs (if present) are to the right of their associated peaks. Reorientation simplifies the subsequent steps in the algorithm. The algorithm then uses a process of successive refinement to identify the largest peaks and then refine the partition to include smaller peaks. Finally, there is a clean-up phase that removes extraneous peaks. The following subsections describe these steps in more detail.

#### 2.2.1 Determining a consensus direction

Because large artifacts can obscure the true direction of the heartbeats, EEG-Beats uses a consensus algorithm to determine how to reorient the signal if necessary. By default, the algorithm divides up the signal into two-second (*flipIntervalSeconds*) intervals and finds the direction of the maximum absolute value in each interval. Only intervals whose most extreme value is more than 1.5 (*threshold*) robust standard deviations from the overall median are considered when determining whether to negate the signal, based on the dominant direction of the extreme points. A similar algorithm determines whether, assuming that a peak is associated with a trough, the trough is before or after the peak. If necessary, EEG-Beats flips/reverses the signal so that most of the peaks are up with following troughs.

#### 2.2.2 Selecting eligible peaks

To determine whether a particular point could really be a heartbeat, the algorithm assumes that eligible points must have a maximum amplitude of at least 1.5 (*threshold*) robust standard deviations from the median and must be at least 500 (*rrMinMs*) ms away from points that have already been determined to be heartbeats. The *rrMinMs* and *rrMaxMs* parameters determine the allowed range of the RR interval sizes. By default these are set to 500 ms and 1500 ms, corresponding to heart rates of 120 bpm and 40 bpm, respectively. While appropriate for normal EEG recordings, these settings should be adjusted for recordings during which the subject is performing strenuous or stressful activities.

In addition to position and amplitude criteria, EEG-Beats uses a sharpness to determine whether a point is an eligible peak. In the case of single peak, the signal at an eligible peak must fall below the median signal within 100 (0.5\**qrsDuration*) ms on either side of the peak value. In the case of a peak followed by a trough, an eligible peak’s trough (as determined by MATLAB’s *findpeaks* function) must be within 100 (0.5\**qrsDuration*) ms of the peak. These checks determine whether a large deviation in signal amplitude actually corresponds to a sharp peak. EEG-Beats applies the algorithm twice (once assuming all peaks are single and once assuming all peaks have following troughs) and then combines the results as described below.

#### 2.2.3 Successively subdividing to find the peaks

The main algorithm uses a divide-and-conquer strategy similar to that used in binary search. EEG-Beats divides the signal into 31 (*consensusIntervals*) equal-size intervals and finds the maximum point (the fence post) in each interval. After adding the first point and the last point in the signal as outer fence posts, EEG-Beats eliminates internal fence posts that are not eligible peaks before beginning divide and conquer.

The divide-and-conquer phase proceeds as follows. For each pair of consecutive fence posts, the algorithm finds the maximum between the first two fence posts in the list. If this maximum point is an eligible peak, the algorithm inserts the point as a fence post between the original two fence posts. Regardless of whether the point is an eligible peak, the algorithm zeros out the signal within a 200 (*qrsDuration*) ms window around this point and continues the process. When there are no eligible peaks between the first two fence posts, the algorithm removes the first fence post and repeats the process with the next pair, until no fence posts pairs remain to be processed.

#### 2.2.4 Cleaning up

Since the signal was truncated, the actual peaks may be slightly off. EEG-Beats makes a pass through the peak list to adjust the actual peak positions to their true maximum positions and then combines the peaks from the two methods (single-peak versus peak-trough). Most peaks from the two methods will be coincident. For unmatched peaks that are within 100 (0.5\**qrsDuration*) ms of each other, EEG-Beats adjusts the position of the peak with the smaller amplitude to be that of the peak with the larger.

A second clean-up step is to remove extraneous peaks by determining whether removing the peak produces a more consistent representation. For that to be the case, a peak’s neighbors must be within 1500 (*rrMaxMs*) ms and the interbeat interval with peak removed is closer to the median interbeat interval than the largest of the interbeat intervals on either side of that peak. EEG-Beats also marks outlier peaks whose absolute amplitude value is either less than 0.5 (*minPeakAmpRatio*) or greater than 2 (*maxPeakAmpRatio*) times the absolute value of the median peak amplitude.

#### 2.2.5 Combining peaks

The final stage uses successive combination to get a single representation of heartbeat. EEG-Beats sets the group (single-peak or peak-trough) with the greatest number of peaks as the base representation and considers the remaining peaks one at a time starting from left to right. EEG-Beats adds the peaks to the base representation as long as they aren’t closer than 500 (*rrMinMs*) ms to peaks in the existing representation. Peaks before the first peak and after the last peak in the base representation are handled slightly differently, with new peaks added from inside outward. EEG-Beats also reports, but does not remove peaks whose amplitudes are likely to be too high or too low to be beats using the standard robust outlier criteria for boxplots. Peaks whose amplitude satisfies the robust outlier criteria are reported as high or low amplitude peaks.

#### 2.2.6 Saving the results

EEG-beats has an EEGLAB plug-in that allows users to run the analysis on single recordings. However, EEG-Beats is meant to be run using scripts on an entire study in a directory tree. Example scripts show how to provide the source root directory for a study and the root directory for saving the results. The script automatically processes all of the EEG .*set* files in the source directory tree and produces plots for each recording similar to those shown in Fig. 1. EEG-Beats also saves a structure (*ekgPeaks*) containing the peaks found for each recording. This structure can be used as input to produce and save a structure containing the RR measures (*rrInfo*) calculated from the interbeat intervals for the entire study. EEG-Beats also provides scripts to do automated analysis of variance and visualizations of RR measure distributions if the user provides a suitably formatted metadata file as described below.

### 2.3 Available RR measures for HRV

In clinical settings, HRV is often analyzed for short-term variations in time windows ranging from 30 seconds to 5 minutes. Long-term variations are often examined for time periods on the order of many hours. The values of several common RR measures depend on the length of the signal, and the computations of other measures are made under the assumption of stationarity. EEG-Beats automated RR measure calculations include both values computed over the entire recording and values calculated over 5 (*rrBlockMinutes*) minute blocks, as recommended by established guidelines (Task Force of the European Society of Cardiology and the North American Society of Pacing and Electrophysiology, 1996). The block offset for successive blocks is set to 0.5 (*rrBlockStepMinutes*) minutes by default. Table 1 lists the RR measures available currently available in EEG-Beats. Spectral ranges are settable. The choice of spectral method heavily influences computed measure values. By default, EEG-Beats uses MATLAB’s *plomb* function to accommodate the uneven spacing of the RR intervals. A preliminary version using short-term Fourier analysis is also available, but not recommended.

**Table 1.** RR measures computed by EEG-Beats.

| Measure | Description | Units | Typical Values |  |
| --- | --- | --- | --- | --- |
| HR | Mean heart rate | bpm | 60–100 (resting) |  |
| RR | Mean interbeat interval | ms | 926±90* |  |
| SDNN | Standard deviation of interbeat intervals | ms | 50±16* |  |
| SDSD | Standard deviation of differences between adjacent RR intervals | ms |  |  |
| rMSSD | Root mean square of successive differences of interbeat intervals | ms | 42±16* |  |
| pNN50 | Fraction of successive interbeat intervals that differ by > 50 ms | – |  |  |
| NN50 | Number of successive interbeat intervals that differ by > 50 ms | – |  |  |
| Power | Area under power spectral curve | ms<sup>2</sup> | 3466±1018<sup>†</sup> |  |
| VLF | Power of interbeat-intervals in very low (0.003-0.04 Hz) range | ms<sup>2</sup> |  |  |
| LF | Power of interbeat-intervals in low (0.04-0.15 Hz) range | ms<sup>2</sup> | 519±291* | 1170±416<sup>†</sup> |
| LFnu | Normalized power of interbeat-intervals in (0.04-0.15 Hz) range | nu | 52±10* | 54±4<sup>†</sup> |
| HF | Power of interbeat-interval spectrum in (0.15-0.4 Hz) range | ms<sup>2</sup> | 657±777* | 975±203<sup>†</sup> |
| HFnu | Normalized power of interbeat-intervals in (0.15-0.4 Hz) range | nu | 40±10* | 29±3<sup>†</sup> |
| LF/HF | Ratio of LF power to HF power | – | 2.8±2.6* | [1.5, 2.0]<sup>†</sup> |

\*Typical values taken from (Nunan et al., 2010). <sup>†</sup>Typical values taken from the Task Force (1996).

**Table 2.**
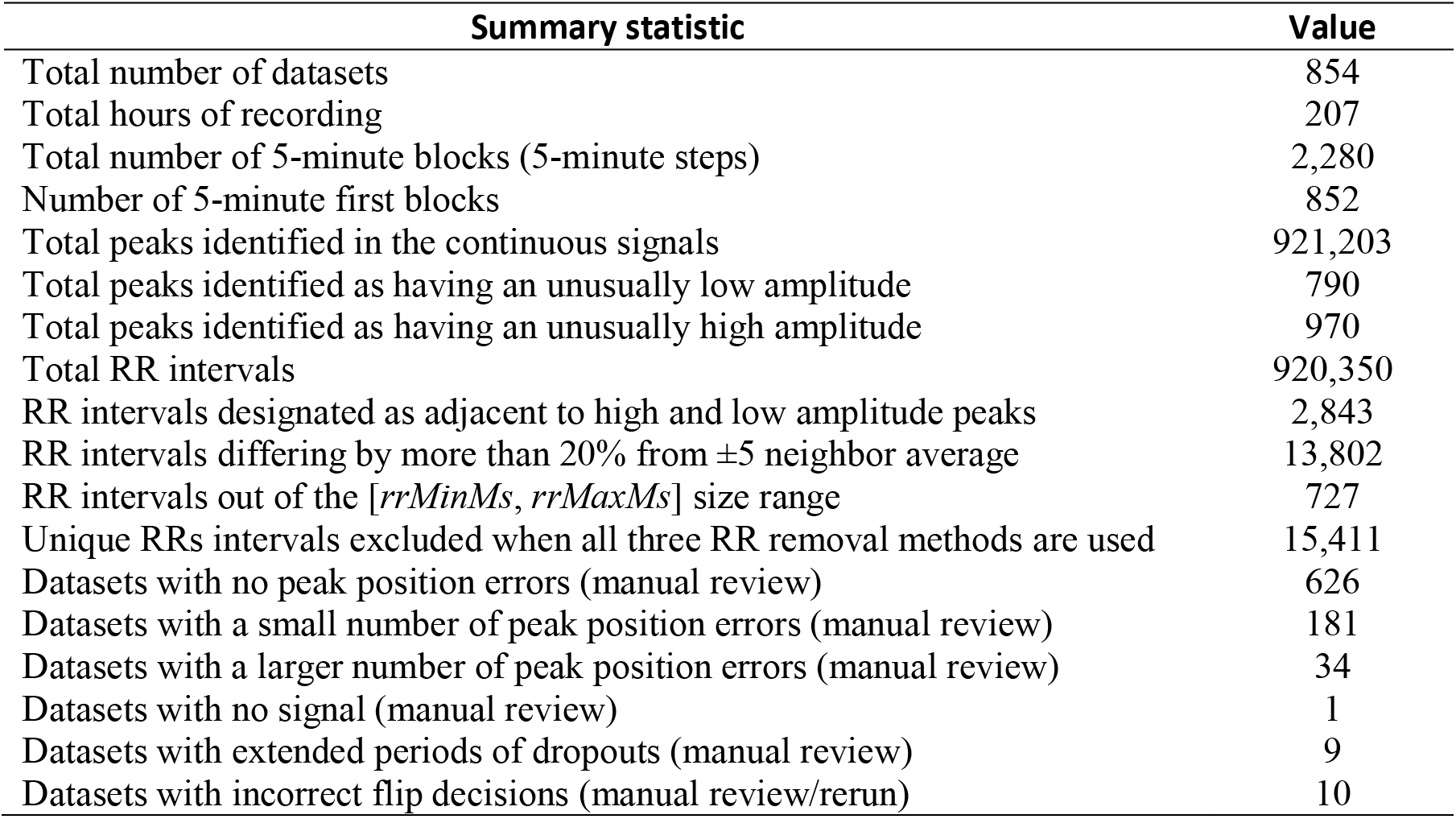
EEG-Beats peak-finding summary for NCTU RWN_VDE.

Another issue that strongly influences the spectral results is the selection of signal detrending method prior to computation of spectral measures. By default, EEG-Beats uses polynomial detrending of order 3 (*detrendOrder).* Total power and low frequency power indicators are higher with no detrending, but power ratios and power in normalized units (LFnu and HFnu) are not heavily influenced by this choice. Normalized units are expressed in terms of the fraction of power in the low frequency range and above.

RR measures are typically calculated after “outlier RRs” are removed. EEG-Beats provides three options for removing outlier RRs:

1. *Outlier RR amplitudes:* Remove RRs outside of [500, 1500] ([*rrMinMs*, *rrMaxMs*]) ms if *removeOutOfRangeRRs* is true.
2. *Outlier peak amplitudes*: Remove one (*rrsAroundOutlierAmpPeaks*) RR interval on either side of peaks with unusually high or low amplitudes,
3. *Bad neighbor RRs:* Remove RRs that differ by more than 20 (*rrPercentToBeOutlier*) percent from the mean of the 5 (*rrOutlierNeighborhood*) neighbors on either side of that RR interval.

Users can choose any combination of three outlier methods for the RR measures calculation. By default, all of three types of outliers are removed.

### 2.4 Analysis and visualization

In addition the visualization of the peak locations and distribution of peaks versus RR intervals (e.g. Fig. 1), EEG-Beats provides a visualization of the RR intervals as a function of time overlaid on (e.g., Fig. 2) or as a subplot with the EKG signal and peaks (Fig. 4). RR intervals detected as outliers using various strategies are marked on these RR interval graphs using different symbols.

**Fig. 2.**
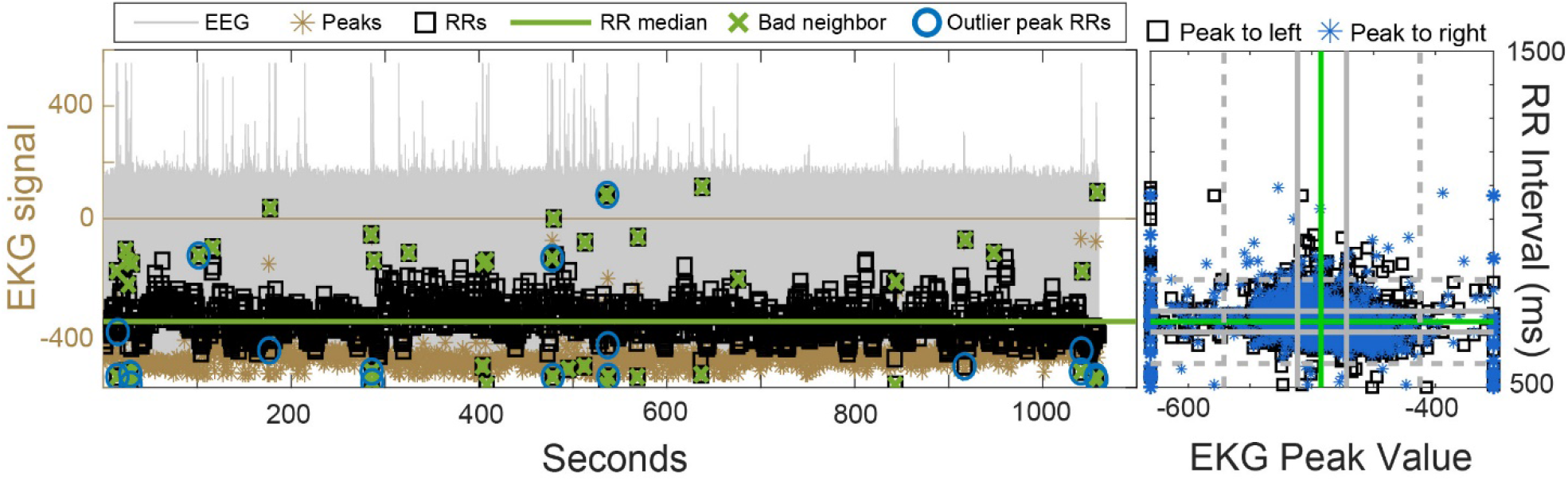
Dataset 800 (subject S21) of the NCTU RWN_VDE study. Left: the EEG-Beats overlay plot. The gray background curve (axis on the left) shows the EKG signal with the peaks shown as gold asterisks. The RR intervals are plotted as a function of time using black squares (axis to the right). The green line shows the median RR interval. Green ×’s mark RR intervals identified as “bad neighbors”, while the blue circles mark RR intervals with outlier amplitudes using the robust outlier criteria. Right: EKG peak amplitude versus RR interval for the peaks on the left (black square) and right (blue asterisks) ends of the RR interval, respectively. Green lines mark medians, and gray lines mark the interquartile ranges for each variable. Dashed gray lines indicate robust outlier limits. The amplitude axes are truncated at 3.0×IQR outside IQR.

EEG-Beats also has several summary visualizations and statistical analyses designed to be run on a study as a whole. The user must provide metadata in a MATLAB structure with one row for each dataset. The structure must have a *fileName* field containing the name of the file that the EKG was extracted from. That *fileName* is matched with the *fileName* field in the *ekgPeaks* structure to perform analysis.

EEG-Beats can compare peak representations beat-by-beat for two different toolboxes (e.g., Table 3) or for output from EEG-Beats for different peak-finding parameter settings. The EEG-Beats RR measure compare functions allow comparison of results from different packages or from using different EEG-Beats artifact removal settings (e.g., Table 4).

**Table 3.** Comparison of EEG-Beats and PNC peak positions.^†^

| Summary counts | EEG-Beats | PNC |
| --- | --- | --- |
| Datasets with peaks | 853 | 773 |
| Datasets which jointly have peaks | 772 | 772 |
| Datasets which jointly have peaks and no drop-outs | 764 | 764 |
| Detected peaks* | 842,079 | 832,286 |
| Peaks with exact frame agreement | 764,657 | 764,657 |
| Peaks unmatched in other | 77,422 | 67,629 |
| Closest peak in other 1 frame away | 19,114 | 19,114 |
| Closest peak in other 2 frame away | 9,951 | 9,951 |
| Closest peak in other 3 frame away | 3,873 | 3,873 |
| Closest peak in other 4 or more frames away | 44,484 | 34,691 |

<sup>†</sup>One frame or sample is equivalent to 7.8 ms of separation. Datasets with dropouts have been removed.

EEG-Beats allows users to form feature vectors representing HRV in each segment (5 minutes by default) by selecting a list of the RR measures from those available from Table 1. EEG-beats normalizes the features by z-scoring and uses MATLAB’s *tsne* function (*t*-distributed stochastic neighbor embedding) to project these high-dimensional feature vectors into a 2D or 3D space (Van Der Maaten & Hinton, 2008). The idea behind *t-SNE* is that high-dimensional vectors projecting to the same low-dimensional clusters are likely to also be similar in the original high-dimensional space. EEG-Beats provides the ability to plot these projections using colors and/or shapes to distinguish different metadata values.

EEG-Beats can also plot the boxplots of any RR measure of Table 1 segregated by any of the metadata variables (e.g., Fig. 6). Finally EEG-Beats can apply an analysis of variance using MATLAB’s *anovan* function for any RR measure using factors corresponding to any combination of fields in the user-provided metadata structure (e.g., Tables 5 and 6).

### 2.5 Data used for testing

This study uses the NCTU RWN_VDE data collection from a longitudinal experiment conducted at National Chiao Tung University (NCTU) during the 2014-15 school year to assess the effects of fatigue and stress on performance (Lin et al., 2016) (Lin et al., 2018) (Huang et al., 2019). Each of the 17 study subjects wore an Actigraph activity monitor for the duration of the study and completed several subjective assessments of stress, fatigue, and sleep quality on a daily basis. Central to the study was the use of the Daily Sampling System (DSS), which processed this information and automatically invited subjects to the laboratory to undergo the experiment based on putative subject state.

The 17 subjects were recorded on up to 9 days, ideally representing low, normal, and high fatigue levels as determined by the DSS. An experimental session consisted of six tasks: resting with eyes open (5 minutes), PVT: psychomotor vigilance (10 minutes), LK: lane-keeping (15 minutes), DAS: driving with distractions presented at a low and high rates counter-balanced (30 minutes each), and concluding with resting eyes open (5 minutes). Each experiment recorded 62 channels of EEG and one channel of EKG using a Neuroscan headset. During each session, the subjects answered several questionnaires before and after each phase of the experiment. Our analysis uses the EKG from the EEG/EKG recordings and the output of the DSS system as a fatigue factor.

## 3 Results

This section presents a validation of EEG-Beats results, a comparison of EEG-Beats with the PhysioNet Cardiovascular Signal Toolbox (PNC), and examples of EEG-Beats analysis tools.

### 3.1 Manual verification of heart beat locations

To validate EEG-Beats peak-finding, we manually reviewed the EEG-Beats peak positions for all 854 EEG data recordings of the NCTU RWN_VDE study using the pan and zoom features for MATLAB figures with the peak and point plots of Fig. 1. A summary is shown in Table 2.

Total peaks indicates the total number of peaks detected in the data corpus. EEG-Beats accurately detected peaks using the default parameter settings in a large portion (626) of the recordings, many of which had substantial artifacts including dramatic variations in amplitude, dropouts, and high-amplitude bursts. Peaks identified as having too high or too low of an amplitude use the standard robust outlier criteria described in the methods. Another 181 datasets had a few errors, while 34 datasets had a “larger” number of errors. Fig. 2 uses an overlay plot to display Dataset 800, an example of a dataset with a “larger” number of errors.

Dataset 800 has a very noisy signal with a number of very low and very high peak amplitudes as indicated by the large number of points on the vertical axes in the right graph of Fig. 2. Although EEG-Beats misidentifies some peak positions for this dataset, all of the misidentifications occur for peaks on one side or the other RR intervals marked either as bad neighbor RRs or as outlier peak RRs on the left graph. Using the three RR outlier removal criteria removes most of the RR outliers in datasets with peak errors.

Most of EEG-Beats misidentifications occur when a peak is overlaid with or is located close to an artifact as illustrated by the example in Fig. 3. The vertical arrow at 620 seconds in the left graph of Fig. 3 marks a large-amplitude artifact that EEG-Beats labeled as a peak. Once that peak was marked as an eligible peak, it became a fencepost during peak detection, and the correct peaks to the right and left were too close to be detected in successive refinement.

**Fig. 3.**
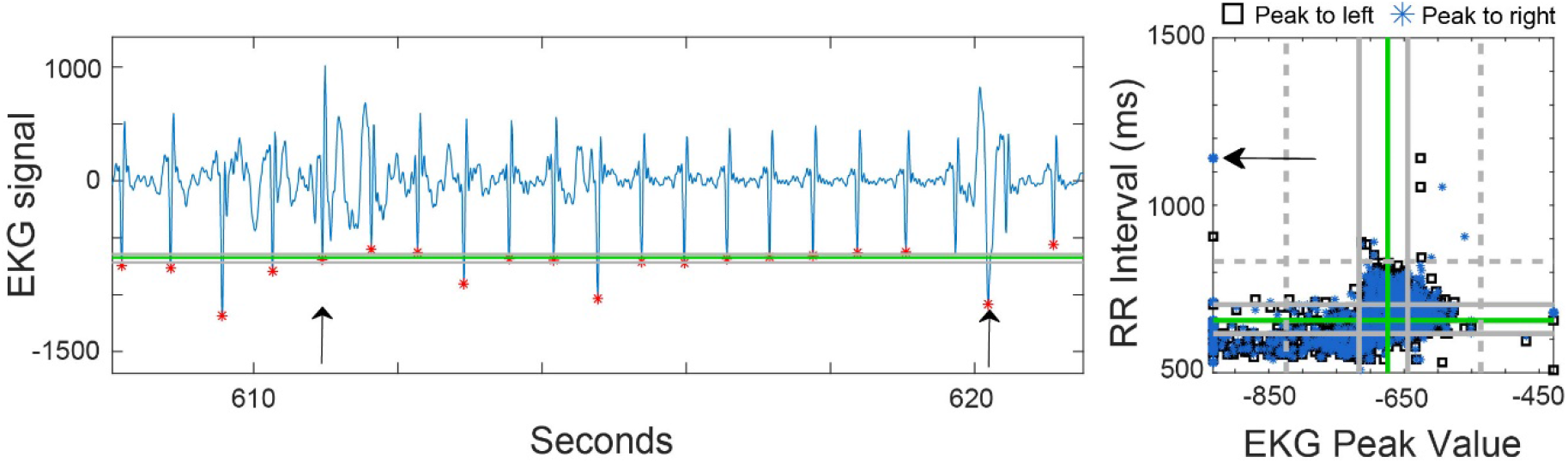
Excerpt from Dataset 128 (Subject S03) of the NCTU RWN_VDE study illustrating problematic peaks. Left: EKG signal acquired using an EEG sensor (blue) with red asterisks indicating the peaks identified by EEG-Beats. The vertical arrow at 611 seconds marks a peak labeled as an ambiguous in-range peak during manual checks. The vertical arrow around 620 seconds marks a peak labeled as incorrect. The two adjacent peaks were marked as missed peaks during manual evaluation. Right: EKG peak amplitude versus RR interval for the peaks on the left (black square) and right (blue asterisks) ends of the RR interval, respectively. Green lines mark medians, and gray lines mark the mid quartile boundaries. Dashed gray lines on the right graph indicate the robust outlier limits. The amplitude axis is truncated at 3.0×IQR outside of the mid quartiles. The horizontal arrow marks the point corresponding to the peak marked at 620 seconds in the left graph.

The right graph of Fig. 3 plots peak amplitude versus the length of the RR intervals immediately to the left (blue asterisk) and to the right (block square) of the peak. This graph, which provides a connection between peaks and RR intervals, can often provide a quick summary of the issues that a given dataset might have. If the plot forms an oval shape with many RR interval lengths falling close to the central quartiles (solid gray horizontal lines), the data is well-behaved. The dashed gray lines indicate the robust outlier thresholds, the traditional thresholds for marking outliers on box plots. This particular dataset has a lot of high negative amplitude peaks falling outside the dashed gray lines, indicating the possible presence of high-amplitude signal artifacts. By default, EEG-Beats clips the amplitude on these plots at 3.0×IQR outside the mid quartiles, so points clustered along the vertical edges of the graph may represent very large or very small amplitude peaks.

In peak *vs*. RR plots, RR markers for RR intervals adjacent to a given peak fall on the same vertical line. Uniform heart rate signals have peak *vs.* RR plots with closely-spaced, vertically-aligned asterisk-square pairs. Similarly, each RR interval is represented by a horizontally-aligned asterisk-square pair corresponding to the peaks at either end of that interval. Widely separated horizontal pairs indicate a dramatic difference in amplitudes between the bounding peaks of the RR interval.

A tip-off that there might be a problem with the peak at 620 seconds is indicated by the horizontal arrow in the right graph of Fig. 3. The lone blue asterisk represents a large amplitude peak with a very long interbeat interval to its right. Further, that large RR has a large negative amplitude (≤ −1000) peak to the left and a much smaller negative amplitude (≥ −650) peak to the right. As a result of incorrectly selecting the artifact as a peak, the RR interval to the right is much longer than that of its neighbors. Thus this RR interval was correctly caught as invalid under the bad-neighbor criterion.

EEG-Beats is able to correctly identify peaks in the face of many artifacts such as the one that appears near at 611 seconds in the left graph of Fig. 3. This artifact is of lower amplitude than the adjoining heartbeat peaks, so it is not considered. However, EEG-Beats is often able to handle artifacts that are much higher than the adjacent beats, provided they don’t meet the sharpness criteria or fall too closely to a real peak.

EEG-Beats has both overlay and subplot time visualizations of the RR intervals. Fig. 4 shows an example of the subplot visualization for Dataset 11 (Subject S01). The top view shows the RR intervals versus time as in the overlay plot, while the middle graph shows the EKG signal with peaks overlaid. The bottom row has an expanded view of the EKG for a small portion of the signal, as well as the peak amplitude versus RR plot on the right. The vertical arrow in the bottom graph marks an extra, low-amplitude EEG peak adjacent to a very small RR interval. The RR interval is marked both with a blue circle (RR amplitude outlier) and a green cross (RR bad-neighbor).

**Fig. 4.**
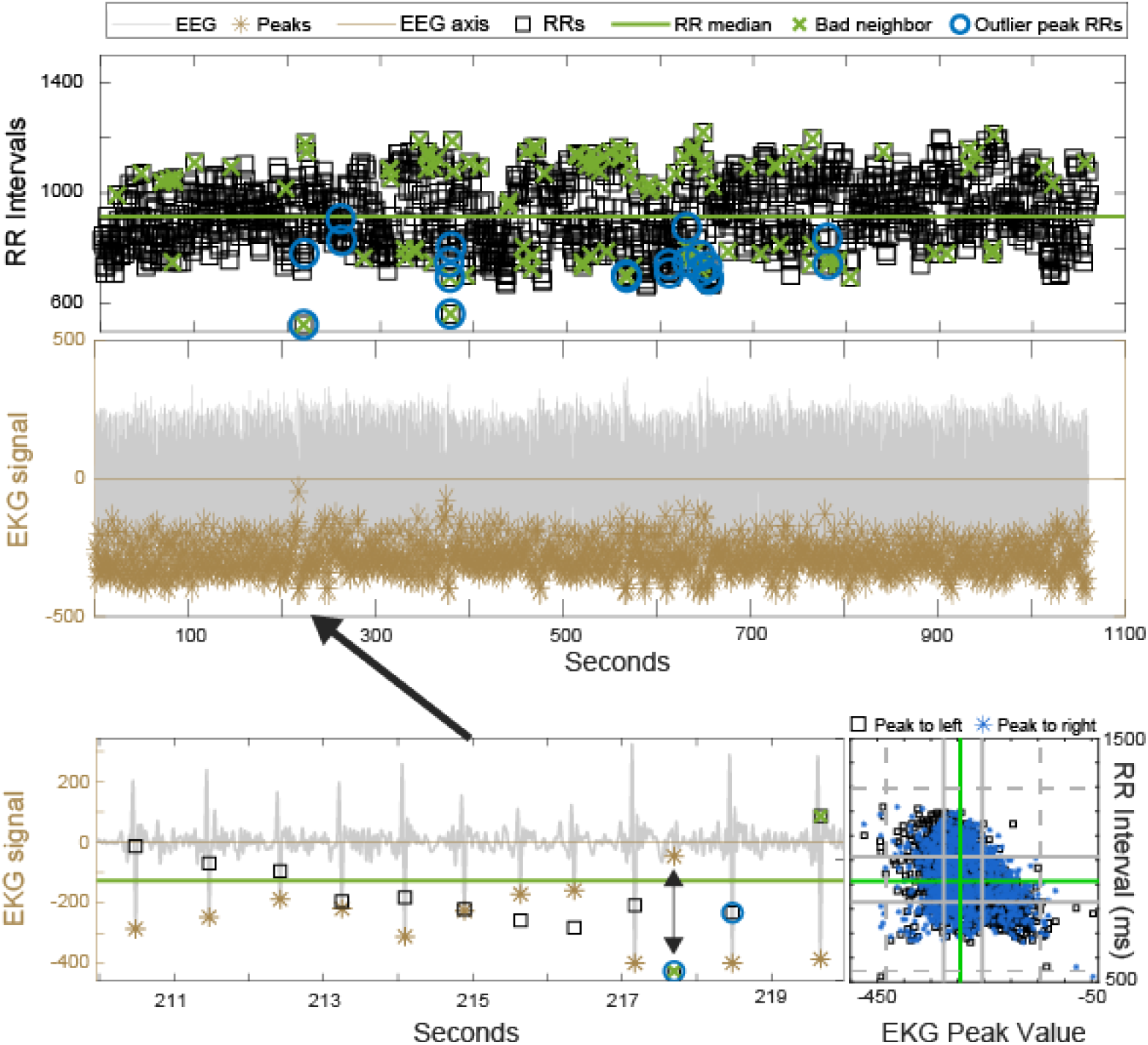
RR interval subplot visualization for Dataset 11 (subject S01) of the NCTU RWN_VDE study (top two graphs). In the top subplot of the visualization, EEG-Beats plots the RR interval lengths versus time as black squares. RRs marked as outliers using the bad neighbor criteria are over-plotted with green ×’s, while RR intervals adjacent to peaks with outlier amplitudes are over-plotted with blue circles. The middle graph plots the EKG signal on the same time line (in gray) with golden asterisks marking the peak positions. The bottom left plot shows an expanded view of a 10-second segment (as indicated by the thick black arrow) using plot overlay form. In this form, the EKG and peaks are plotted using the left axis and the RR interval values and outlier markers use the right axis. The vertical double arrow indicates that the outlier RR with value near 218 seconds is associated with the presence of the extraneous peak. The graph in the lower right corner is the peak versus RR interval graph for the dataset. The extraneous peak appears in the lower right graph as a blue asterisk in the lower left corner (small amplitude peak to the right of a very small RR interval).

On the surface this dataset falls into the “larger number of incorrect peaks” category as the dataset in Fig. 2. In fact, this dataset just had a few incorrectly identified peaks — all of which were associated with outlier RR amplitudes. The “bad-neighbor” criteria had many “false positives”. While many of the subjects in the study have slowly varying interbeat intervals, some subjects such as Subject S01 had heart rates that were quite variable over very short periods of time. This short-time scale variability is quite clearly visible in the inset shown in the bottom row of Fig. 4.

Unlike the example of Fig. 2, where the green crosses were clearly outliers from a slowly varying RR interval amplitude trend, the green crosses in this case are not clearly distinguishable as outliers. The bad neighbor criteria for eliminating outlier RRs works well for subjects with heart rates that are relatively steady or are slowly varying over time scales larger than the “neighborhood”. However, it is not clear how useful or applicable such a criterion is when analyzing individuals with more unusual heart rate patterns.

Nine of the datasets had extended periods of dropouts. An example appears in Fig. 5 for Dataset 587 (Subject S17). The RR artifact detection works well to remove outliers, although this dataset should be trimmed before using in analysis. The peak vs. RR interval amplitude shown in the right graph clearly exposes the abnormality, as the points are aligned vertically at large amplitudes. The vertical line to the right would expand into a normally shaped ovoid if the signal before 100 seconds were removed from the data record.

**Fig. 5.**
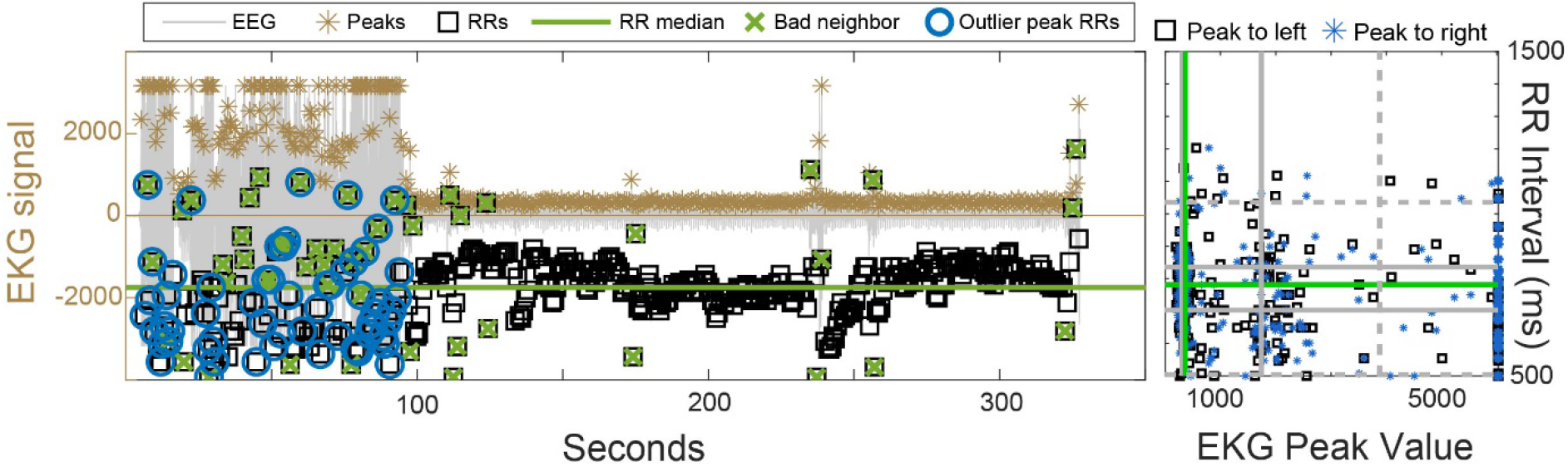
RR interval overlay visualization for Dataset 587 (subject S17) of the NCTU RWN_VDE study. Caption as in Fig. 2. This dataset was one of the nine datasets identified as having extended periods of dropouts.

**Fig. 6.**
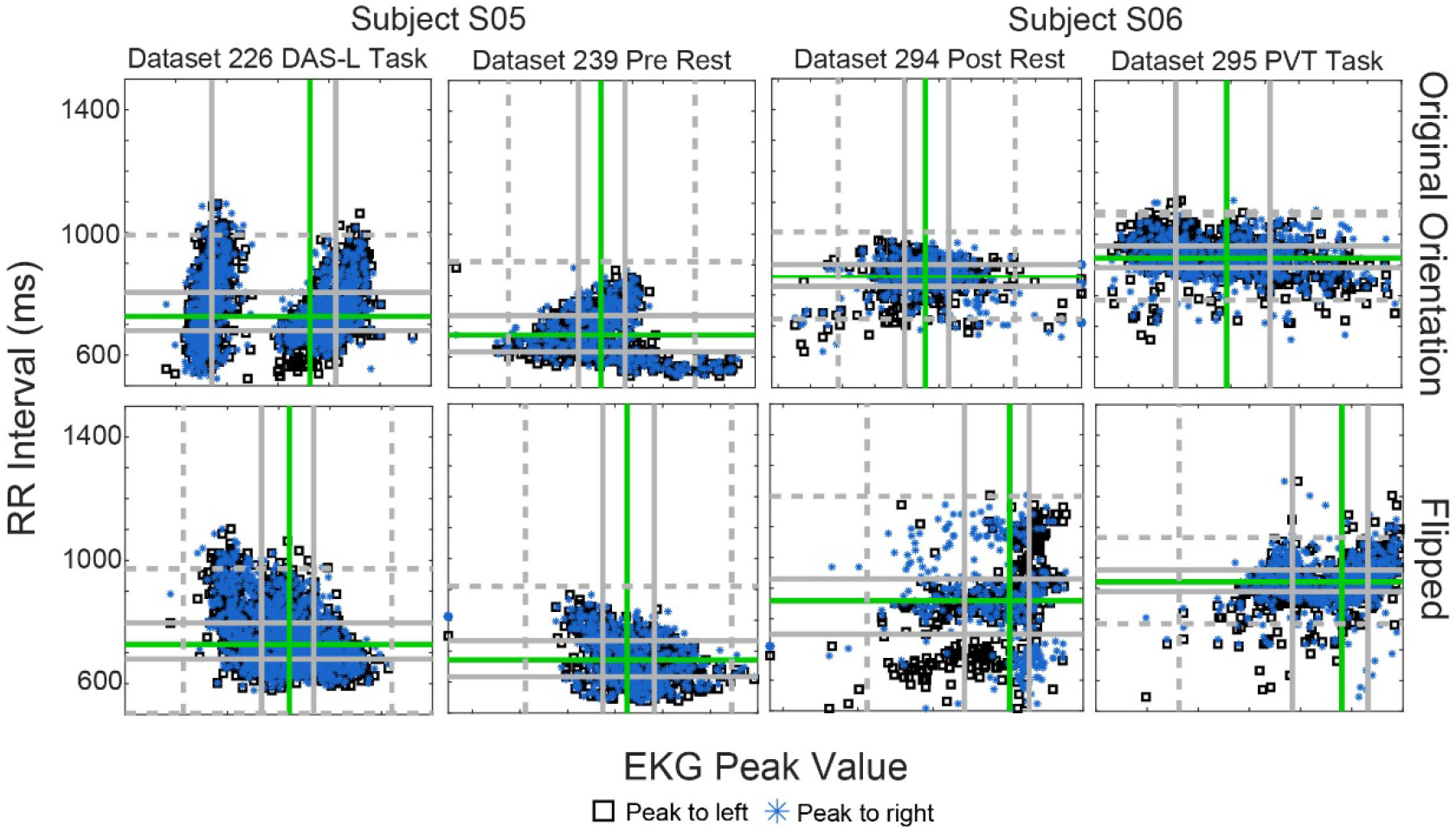
EKG peak amplitudes vs. RR interval to the left (blue asterisk) or to the right (black square) for four datasets from two subjects acquired on different days. Each column corresponds to the same data set in original orientation (top graph) and flipped orientation (bottom graph). (See Fig. 1 caption for additional explanation).

A final category of problematic datasets listed in Table 2 are the ten datasets in which EEG-Beats picked the wrong direction for the peak orientation (either flipping when it shouldn’t have or not flipping when it should have). Flipping errors are usually not apparent from the EKG signal, but show up in the non-ovoid shapes of peak-interval plots as illustrated in Fig. 6.

Dataset 226 from Subject S05 (first column of Fig. 6) had a very intermittent EKG signal, switching at random between unidirectional peaks with small troughs and peak-troughs with larger troughs. The two sets of amplitudes correspond to sections with small troughs and larger troughs, respectively. With the flipped signal EEG-Beats detected peaks centered on an intermediate amplitude. The actual RR interval distribution changed very little since the either choice gave a consistent peak spacing, but the correct orientation resulted in more accurate peak detections. (With the correct orientation, the dataset moved from having a larger number to having no errors during manual verification.) The second column shows the results from Dataset 339, also from Subject S05 but from a different session. This dataset had peaks and troughs of roughly equal amplitude. It turned out that the amplitude of the “peaks” was more variable than that of the troughs, resulting in the long tail in amplitude. When flipped that dataset had no peak detection errors, while in the original orientation, it had a few errors on manual review.

The third column of Fig. 6 (Dataset 294 Subject S06) shows an example of the third class of issues EEG-Beats encountered in making a flip decision. The peaks in these datasets looked like three-lobe triplets with almost equally deep troughs. The dominant amplitude within the triplets in the direction EEG-Beats picked as up sometimes changed, resulting in three different spacing groups (hence three horizontal lines in the plot). It happened that the dominant amplitude in the troughs was consistently the central trough, so in that orientation gave a better distribution. The fourth column of Fig. 6 (Dataset 295 Subject S06) has the same alternation between unidirectional and peak-trough during the recording. Here the two clusters are less prominent because the differences in amplitude aren’t as great.

Several observations can be made about the graphs in Fig. 6. Since these issues did not appear in recordings from other sessions with these subjects, we can conclude that the issues occurred because of improper seating or electrode replacement. Notice that regardless of the orientation, EEG-Beats was able to provide a fairly consistent RR distribution. The two columns associated with each subject correspond to recordings made on different days — one with an active task and the other with a resting task. Heart rates appeared to be lower in the resting tasks for both subjects.

### 3.2 Comparison of heart beat detection with the PNC toolbox

As a point of reference, we compared peak locations determined by EEG-Beats and the PhysioNet Cardiovascular Signal Toolbox (PNC) for the first 5-minute block in each dataset. When used with its default parameters, PNC failed on almost all datasets due to rejection because of atrial fibrillation detection. The results in this section bypassed PNC’s atrial fibrillation detection.

Table 3 compares peak detection results from EEG-Beats and PNC using the default parameters for both toolboxes. EEG-Beats and PNC detect peaks at exactly the same places in most cases. The PNC predictions that are 1 to 3 frames off from EEG-Beats are not located at the exact frame positions of the signal maxima, which can happen when amplitudes are thresholded. EEG-Beats compensates for this problem by adjusting the peaks to the exact peak maxima in the unthresholded signals at the end of peak finding. EEG-Beats detects beats in all datasets but one that it correctly identifies as containing just noise. PNC detect beats in this non-signal dataset, but was unable to detect beats in 81 other datasets. Dataset 11 of Fig. 4 was such a dataset. Most traditional beat detection algorithms are not prepared to deal with the artifacts and the varying peak amplitudes that are common to EEG monitoring of EKG. The peak matches were computed after the ten datasets that were found to have drop-outs were removed.

### 3.3 Comparison of RR measures with the PNC toolbox

PNC also allows users to provide the RR intervals instead of raw EEG signals and computes an extensive collection of RR measures. Since PNC is well-benchmarked, we provided the RR intervals computed by EEG-Beats to PNC and computed RR measures for comparison with EEG-Beats. The results are summarized in Table 4. The values are based on the first 5-minute block in each of the 853 datasets for which there was a signal. PNC was not able to RR measures on 219 of these datasets. We attribute the rejection by PNC to certain internal rejection procedures it applies when calculating the measures.

The default RR artifact removal was used for both toolboxes. For EEG-Beats, this means that out-of-range RRs are removed as well as bad neighbors and the RRs on either side of peaks with unusually high or low amplitudes. Exclusion of these RRs changes the results by a small amount. For comparison, the EEG-Beat calculation on the raw RRs (no removal) are shown in parentheses. The differences between the means of the two values are not large. For most, but not all, measures the agreement between PNC and EEG-Beats with artifact removal is better than with raw EEG-Beats values as indicated by the mean absolute differences.

**Table 4.**
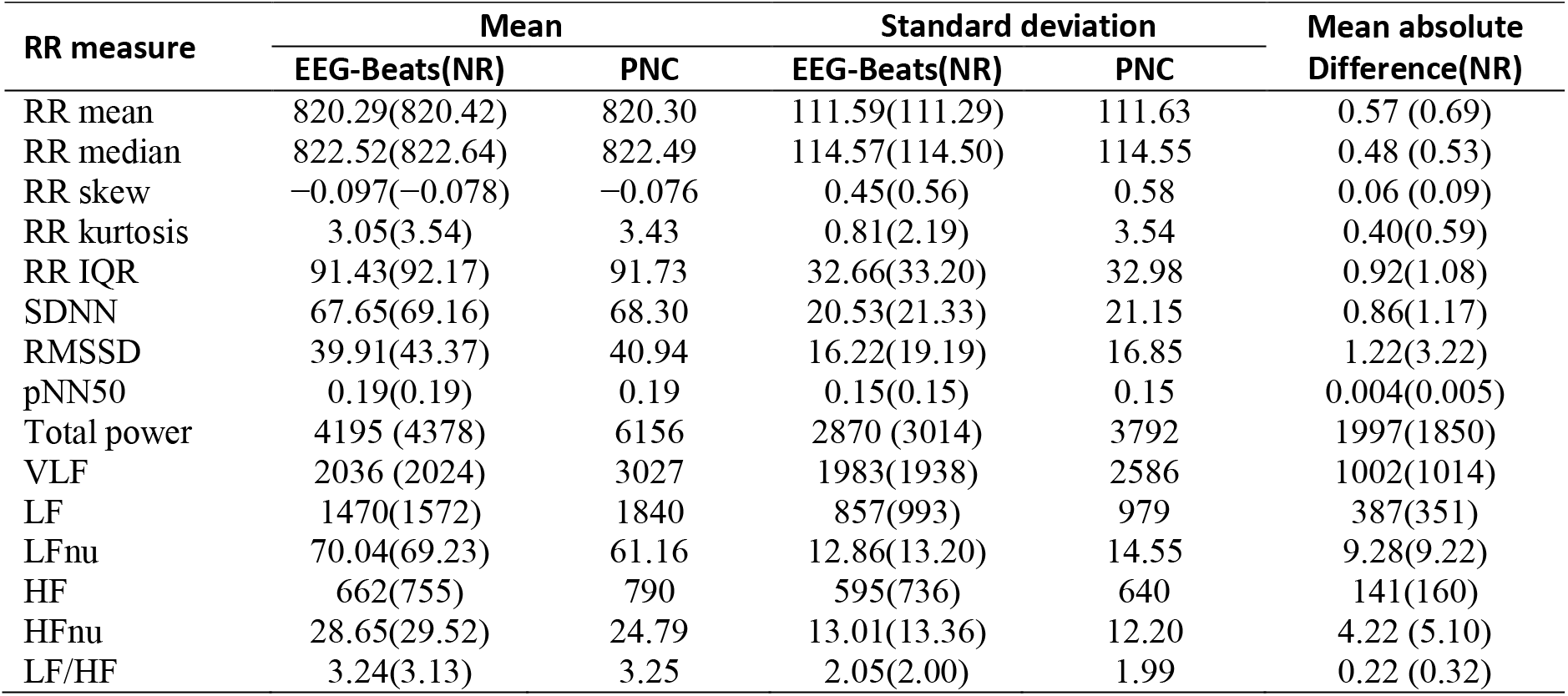
Comparison of EEG-Beats and PNC RR measures. NR=no removal of RR artifacts.

In general, the agreement between the values computed by EEG-Beats and PNC is quite close. Several differences in assumptions may explain some of the differences between the values computed by the two toolboxes. EEG-Beats uses a default interval range for RRs of [500, 1500] ms, while PNC’s default range for RR intervals is [375, 2000] ms. Both algorithms use the same default neighbor value exclusion. By default, both EEG-Beats and PNC use *plomb* to compute the RR interval spectrograms for computation of the frequency measures, however, the assumed frequency resolution differs as does approach to trend removal. PNC removes the mean of the signal prior to spectral computation, while by default EEG-Beats removes a polynomial (cubic) trend, resulting in smaller overall total power and smaller low frequency power for EEG-Beats. However, the non-dimensional power ratios for the two methods are similar. The EEG-Beats trend removal options are settable. The time measure predictions differ slightly, most likely due to the additional RR artifact removal steps applied by PNC.

### 3.4 EEG-Beats visualizations

EEG-Beats provides scripts for creating boxplots of any RR measure segregated by the values of a metadata variable such as subject or task. The user must provide the study metadata in a structure with one line per dataset. The fieldnames of the structure can used to specify how to select and group metadata variables for the boxplots. Fig. 6 shows an example that was generated by EEG-Beats for the LFnu measure with subject IDs as the group variable. All datasets were used with no adjustments.

The variability in the distributions of the unscaled values across subjects (top graph of Fig. 7) is typical of most RR measures. Individuals clearly have baseline values that differ quite widely. EEG-Beats provides two methods of scaling out some of the individual variability: by division or by subtraction. The user must specify a particular baseline block of data for the scaling. In the case of NCTU RWN_VDE, a 5-minute block of resting data acquired at the beginning of each experimental session was an obvious baseline. The scaling reduces variability of the indicators across subjects. However, for the RWN_VDE data collection, the sessions were scheduled at different levels of the nominal subject fatigue level, and scaling by a session-specific value reduces the effect of daily fatigue level on the subject’s measures. A better approach for this data collection might be to scale by the average of the baselines over all of a subject’s sessions, but this is not supported as an automated EEG-Beats feature.

**Fig. 7.**
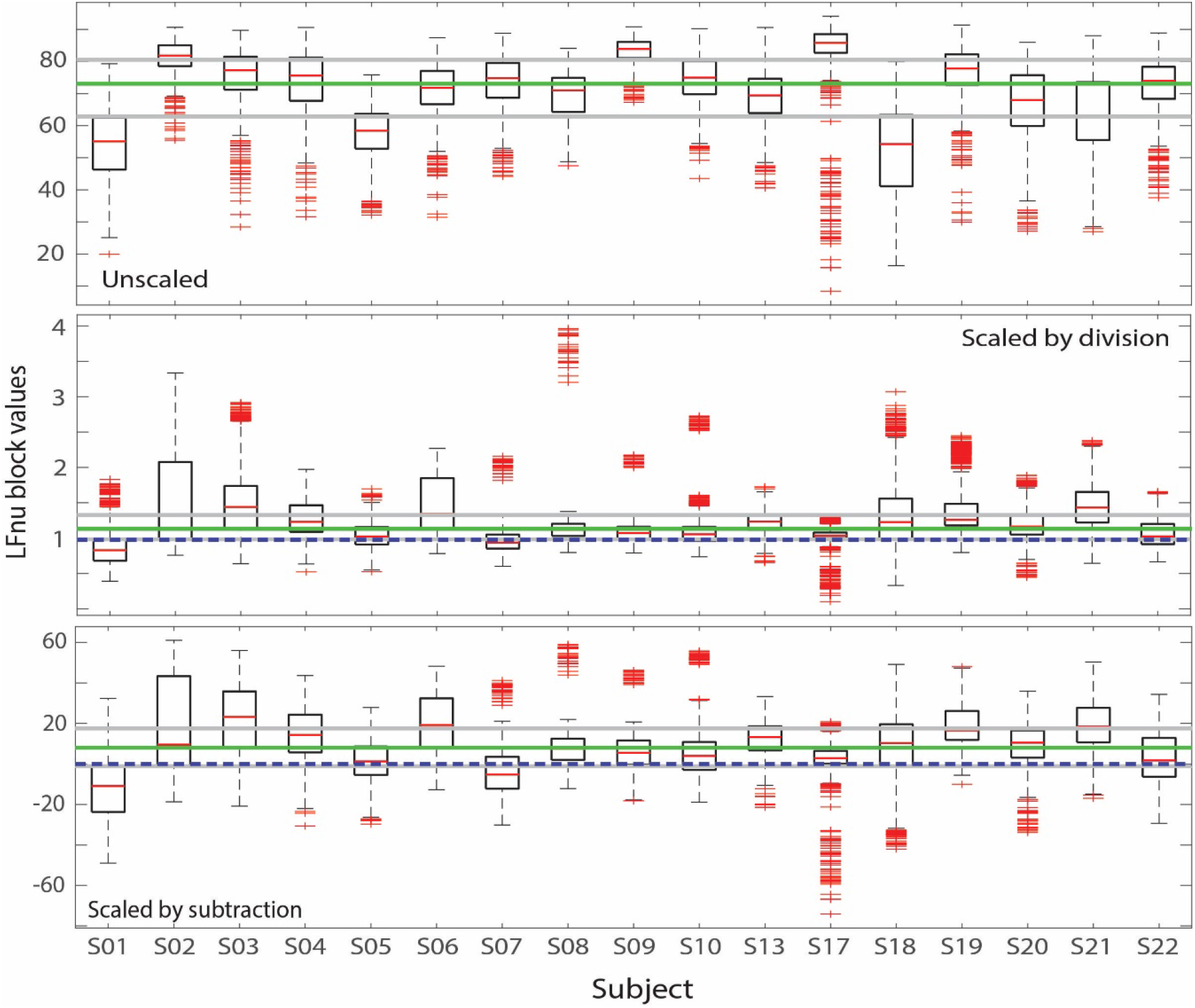
Distribution of LFnu (low frequency power normalized units) segregated by subject as calculated and displayed by EEG-Beats. Top: Unscaled values. Middle: Values scaled by division of a base value. Bottom: Values scaled by subtraction of a base value. See text for explanation of base value determination. Values were computed in 5-minute sliding windows with a 0.5 minute window offset. The plot uses measures computed after RR artifacts were removed. The solid green line marks the overall data median with solid gray lines marking the mid quartile boundaries. The dark blue dashed lines mark the baseline scaled values in their respective plots. The boxplots use MATLAB’s data defaults.

### 3.5 EEG-Beats factor analysis

The EEG-Beats toolbox includes scripts for study-wide analysis of variance (ANOVA). As with the boxplots, EEG-Beats relies a metadata structure provided by the user and can analyze an arbitrary number of pairs of factors identified by the metadata structure fieldnames. The NCTU RWN_VDE data set has subject, task, fatigue level (DSS measure), gender, and replicate number as possible group variables. Table 5 shows an example of the output of ANOVA analysis using task and fatigue levels as factors. Most of the RR measures in Table 5 show a significant statistical dependence on task and on nominal fatigue level at the 0.01 significance level. Significance is usually improved with scaling. The RR measures were computed from data in which RR artifactual values were removed using the default settings.

**Table 5.** P-values for ANOVA for task versus nominal fatigue level. Shaded values are not significant at the 0.01 significance level.

| RR Measure | Unscaled |  |  | Scaled by subtraction |  |  |
| --- | --- | --- | --- | --- | --- | --- |
|  | Task | Fatigue | Task×Fatigue | Task | Fatigue | Task×Fatigue |
| Mean HR | 1.98E-25 | 7.92E-08 | 2.70E-03 | 1.91E-40 | 5.74E-10 | 4.52E-10 |
| Mean RR | 1.13E-27 | 9.20E-10 | 4.65E-04 | 1.38E-38 | 1.82E-11 | 5.51E-08 |
| Median RR | 4.27E-28 | 2.99E-09 | 4.95E-04 | 1.56E-39 | 1.69E-12 | 1.89E-07 |
| Skew RR | 3.92E-51 | 1.10E-01 | 1.42E-06 | 7.21E-48 | 0.00043 | 3.82E-05 |
| Kurtosis RR | 2.12E-21 | 2.29E-02 | 6.19E-10 | 1.54E-04 | 4.34E-04 | 1.95E-07 |
| IQR RR | 1.80E-93 | 1.30E-22 | 1.18E-21 | 4.13E-90 | 5.96E-22 | 4.82E-22 |
| SDNN | 3.01E-100 | 1.38E-26 | 2.41E-17 | 6.46E-101 | 1.38E-25 | 8.15E-16 |
| SDSD | 5.58E-20 | 5.41E-18 | 1.30E-10 | 2.24E-22 | 1.80E-09 | 4.26E-12 |
| RMSSD | 5.05E-16 | 4.01E-15 | 5.61E-09 | 4.47E-20 | 1.72E-07 | 6.84E-10 |
| NN50 | 6.16E-04 | 1.83E-11 | 1.47E-06 | 6.03E-12 | 1.79E-05 | 1.07E-09 |
| pNN50 | 4.41E-10 | 1.80E-14 | 1.17E-07 | 2.36E-18 | 3.41E-07 | 6.49E-09 |
| Total Power | 2.82E-26 | 9.46E-07 | 1.09E-10 | 1.48E-25 | 2.52E-04 | 6.38E-12 |
| VLF | 1.29E-65 | 4.01E-17 | 1.30E-12 | 1.23E-64 | 7.05E-15 | 8.20E-13 |
| LF | 1.60E-32 | 1.24E-07 | 1.62E-12 | 5.65E-33 | 2.62E-06 | 8.35E-13 |
| LFnu | 1.41E-66 | 4.60E-03 | 4.02E-10 | 5.70E-50 | 2.16E-11 | 6.27E-07 |
| HF | 2.31E-08 | 1.79E-02 | 2.63E-09 | 4.92E-09 | 6.07E-02 | 7.48E-11 |
| HFnu | 1.69E-62 | 4.70E-03 | 7.31E-10 | 3.17E-52 | 3.38E-12 | 1.24E-06 |
| LF/HF | 4.13E-39 | 6.56E-04 | 1.89E-05 | 1.24E-42 | 1.13E-04 | 3.74E-07 |

Caution should be exercised when interpreting these results. Subject-task analysis of variance also showed highly-significant dependence on those factors. However, when subject-fatigue two-way analysis of variance was performed, subject dependence was found to be very highly significant, but fatigue level was not. The interaction between the two was significant, however. This result has a physical explanation. In this experiment, subjects were invited into the lab when the results of the daily sampling system indicated that they were in a particular fatigue state. How these measurements reflect on performance is highly individualized, so the fatigue-level designations are not independent of subject.

EEG-Beats also allows three-way analysis of variance. Table 6 shows the results of subject-task-fatigue analysis. Subject and task are highly significant factors for all RR measures, but fatigue has no effect. This points to the importance of analyzing heart rate variability for subjects individually.

**Table 6.** P-values for 3-way ANOVA for subject-task-fatigue. Shaded value is not significant at the 0.01 significance level. Fatigue was not significant for any measure and is not shown.

| RR Measure | Unscaled |  | Subtraction scaling |  | Division scaling |  |
| --- | --- | --- | --- | --- | --- | --- |
|  | Subject | Task | Subject | Task | Subject | Task |
| Mean HR | 0 | 2.86E-65 | 0 | 8.69E-70 | 0 | 6.61E-69 |
| Mean RR | 0 | 1.54E-81 | 0 | 6.28E-73 | 0 | 1.54E-72 |
| Median RR | 0 | 1.51E-82 | 0 | 1.80E-75 | 0 | 7.52E-75 |
| Skew RR | 8.92E-191 | 3.53E-81 | 2.90E-64 | 2.73E-64 | 8.92E-14 | 1.70E-01 |
| Kurtosis RR | 5.27E-47 | 5.88E-37 | 3.88E-106 | 2.50E-12 | 1.65E-75 | 1.37E-14 |
| IQR RR | 1.08E-143 | 1.90E-161 | 8.77E-168 | 3.30E-155 | 1.37E-260 | 9.58E-101 |
| SDNN | 1.52E-188 | 2.32E-171 | 4.17E-199 | 2.47E-181 | 4.35E-318 | 3.43E-129 |
| SDSD | 0 | 6.56E-44 | 0 | 1.79E-46 | 0 | 81.04E-41 |
| RMSSD | 0 | 2.80E-37 | 0 | 9.52E-42 | 0 | 7.99E-36 |
| NN50 | 0 | 1.04E-16 | 0 | 4.14E-22 | 6.46E-62 | 3.70E-18 |
| pNN50 | 0 | 3.14E-32 | 0 | 4.04E-38 | 9.57E-64 | 3.49E-19 |
| Total Power | 3.29E-13 | 1.64E-30 | 6.18E-29 | 1.00E-30 | 1.11E-47 | 8.84E-16 |
| VLF | 2.03E-46 | 1.35E-96 | 4.22E-19 | 6.16E-103 | 2.16E-93 | 2.19E-59 |
| LF | 9.65E-21 | 6.77E-37 | 2.25E-45 | 2.16E-38 | 1.77E-54 | 3.19E-21 |
| LFnu | 0 | 3.15E-160 | 0 | 3.91E-97 | 1.01E-265 | 1.15E-69 |
| HF | 4.54E-08 | 3.00E-08 | 6.87E-22 | 4.92E-09 | 8.49E-13 | 1.07E-05 |
| HFnu | 0 | 1.54E-164 | 0 | 1.59E-101 | 0 | 2.37E-70 |
| LF/HF | 0 | 2.82E-98 | 7.21E-280 | 8.03E-74 | 4.31E-172 | 3.71E-58 |

The results also show that task is an important factor. In a detailed longitudinal study, Spangler et al. (Spangler et al., 2020) has recently shown that performance and HRV indicators were strongly individualized, but that there were consistent session-level, task-level, and block-level dependencies common across subjects.

## 4 Discussion/Conclusion

No automated algorithm completely survives its first encounter with real data. In large-scale computation, it is important to recognize that even adaptive approaches will fail in cases when the data is bad enough. Further, even when an algorithm works well, a myriad of parameter settings may affect final results. EEG-Beats focuses not only on automating well, but also on providing tools for quickly assessing failures and evaluating dependencies of parameter choices on end results for an entire collection.

EEG-Beats has utilities for beat-by-beat comparisons of two versions of the RR intervals for the same dataset. Scripts provide overall summaries of agreements (such as those produced for Table 3 for the EEG-Beats *vs.* PNC beat comparisons). These scripts can be also used on two versions of the RR intervals produced by EEG-Beats for different values of any parameters to see how much the end results are affected by changes in algorithm parameters. For example, we might try a lower setting of the default 500 ms value of *rrMinMs*, (which corresponds to a heart rate of 120 bpm) to see whether the default setting correctly captures all of subject records. The 500 ms settings may not be appropriate for subjects recorded during vigorous activity. The point is that any suspect parameter setting can be quickly tested and the results summarized, either for individual datasets or for the entire study.

Another example is calculating differences in measure values produced by different algorithms for the same set of RR intervals as illustrated by the RR measure comparison of results for EEG-Beats and PNC as presented in Table 4. Also presented in Table 4 is a comparison of EEG-Beats results with and without its RR interval outlier removal.

The EEG-Beat visualizations are designed to enable researchers to assess quickly whether something has gone wrong for a particular dataset. We use the very-large icon preview in the file browser to quickly spot outliers. The RR interval versus EEG peak amplitude plots are particularly useful for assessing whether a dataset might have issues. Non-ovoid distributions, distributions with multiple clusters, or distributions with long trailing or leading tails merit closer inspection (Fig. 6). Almost all problematic datasets can by spotted by looking at these previews.

The development of EEG-Beats was motivated by the potential for using heart rate variability as a low-cost secondary measure of subject state in EEG experiments. We started looking at EEG for EKG when a careful analysis of interbeat interval information provided as output by the wrist monitors and bioharness detectors also used in these experiments showed inconsistencies and numerous dropouts. We developed EEG-Beats after encountering substantial difficulties in applying standard sliding-window approaches to peak-finding due to EEG artifacts. EEG-Beat’s top-down divide-and-conquer approach to peak-finding is able to handle a variety of difficult artifactual signals in an automated fashion, but it is not applicable for online applications. Further, because EEG-Beats focuses on peak detection rather than detection of QRS complexes, it is not suitable for clinical applications and is designed to be used on recordings from normal subjects.

We have shown good agreement with the well-benchmarked PhysioNet Cardiovascular Signal Toolbox (PNC) for cases in which PNC can detect peaks. EEG-Beats is organized into a peak finding stage (*eeg_beats*) that produces a structure containing detailed peak information and a computational stage (*eeg_ekgstats*) that takes this structure and outputs a structure containing the RR measures. Scripts are provided to run these functions on an entire study in an automated fashion and to perform the analyses demonstrated in this paper. EEG-Beats also has an EEGLAB plugin and an associated GUI is under development. EEG-Beats is freely available at https://github.com/VisLab/EEG-Beats. Structures containing NCTU RWN_VDE EKG signals, metadata, and heartbeats are being released as supplemental material as part of this paper to allow other researchers to compare their algorithms.

## 6 Author Contributions

All of the authors worked on algorithm development and analysis during the course of the study. ST, SM, and KR performed manual data evaluations. KR conceived the study and wrote the initial paper draft. All authors participated in the editing and revision of the manuscript.

## 8 Acknowledgments

The authors would to thank Ching-Teng Lin and Jung-Tai King of the BRC (Brain Research Center) at NCTU (National Chiao-Tung University), Taiwan for allowing the use of their experimental data. The authors also appreciate the perspectives on heart rate variability and its relationship to physiology provided by Scott Kerick and Justin Brooks of the Army Research Laboratory, Aberdeen, MD.

